# Patient mutations in DRP1 perturb synaptic maturation of cortical neurons

**DOI:** 10.1101/2024.08.23.609462

**Authors:** TB Baum, C Bodnya, J Costanzo, V Gama

## Abstract

With the advent of exome sequencing, a growing number of children are being identified with *de novo* loss of function mutations in the dynamin 1 like (*DNM1L)* gene encoding the large GTPase essential for mitochondrial fission, dynamin-related protein 1 (DRP1); these mutations result in severe neurodevelopmental phenotypes, such as developmental delay, optic atrophy, and epileptic encephalopathies. Though it is established that mitochondrial fission is an essential precursor to the rapidly changing metabolic needs of the developing cortex, it is not understood how identified mutations in different domains of DRP1 uniquely disrupt cortical development and synaptic maturation. We leveraged the power of induced pluripotent stem cells (iPSCs) harboring DRP1 mutations in either the GTPase or stalk domains to model early stages of cortical development *in vitro*. High-resolution time-lapse imaging of axonal transport in mutant DRP1 cortical neurons reveals mutation-specific changes in mitochondrial motility of severely hyperfused mitochondrial structures. Transcriptional profiling of mutant DRP1 cortical neurons during maturation also implicates mutation dependent alterations in synaptic development and calcium regulation gene expression. Disruptions in calcium dynamics were confirmed using live functional recordings of 100 DIV (days in vitro) mutant DRP1 cortical neurons. These findings and deficits in pre- and post-synaptic marker colocalization using super resolution microscopy, strongly suggest that altered mitochondrial morphology of DRP1 mutant neurons leads to pathogenic dysregulation of synaptic development and activity.

## Introduction

Efficient, long-range transport of mitochondria is essential for proper cortical development^1,2,3^. This is evidenced by the fact that immature neurons have more highly motile mitochondria than fully mature neurons to meet the metabolic needs of a rapidly changing cellular architecture^4^. Mitochondrial fission, or fragmentation, is a critical precursor to mitochondrial transport within the highly elongated morphology of neurons in order to travel through narrow axons^5,6,7^. The large GTPase dynamin-related protein 1, DRP1, is both a necessary and sufficient executor of mitochondrial fission^8^. Once mitochondria are transported along microtubule highways to the distal, synaptic end of the cell, they join a local mitochondrial network where they can support fluctuating demands of ATP production and calcium buffering to support synaptic function^9,10^.

Advances in whole exome sequencing have identified a growing number of patients with mutations in *DNM1L*, the gene encoding DRP1. These patients display severe neurodevelopmental phenotypes, including optic atrophy, developmental delay, microcephaly, and epileptic encephalopathies; most of these patients do not survive past childhood or early adolescence (**Table 1**)^11,12^. There is therefore a growing need to understand why early cortical development is particularly sensitive to disruptions in mitochondrial fission. Despite the severe nature of highly fused networks of elongated mitochondria in patient cells, it is striking that any of these patients can support cortical networks during the beginning stages of fetal development and survive until birth. This study seeks to better understand how cortical neuron development and synaptic maturation is impacted in cortical cultures with mutations in DRP1.

**Table 1.** Clinical Phenotypes of Patients with Variants in Mitochondrial Dynamics Genes. Patients with mutations in DRP1 display severe symptoms of neurodevelopmental regression, atrophy, and altered mitochondrial morphologies. *Mutation results in a stop codon. **Symptoms reported in single patient, not reported in other patients with variants in this domain.

| Variant | Patient Phenotypes | Mitochondrial Morphology | References | PMIDs |
| --- | --- | --- | --- | --- |
| **DNM1L (DRP1)**<br>**G Domain:** E2A, G32A, S39N, S39G, T59N, W88fs, T115M, S36G/E116Kfs*6, W88Mfs*/E129K*6, D146N, G149R, A192E, K206N, K206E, G223V**, L230dup, R237W | optic atrophy, hypotonia, developmental delay, microcephaly, nystagmus, muscle tone abnormalities, ataxia, encephalopathy, mild hearing loss, seizures** | hyperfilamentous, elongated mitochondria, giant mitochondria, reduced mtDNA content, normal peroxisome morphology | Gerber et al 2017, Whitley et al 2018, Liu et al 2021, Keller et al 2021, Lhuissier et al 2022, Hogarth et al 2018, Yoon et al 2016, Nasca et al 2016, Longo et al 2020, Wei et al 2021, Nolden et al 2022, von Spiczak et al 2017 | 28969390, 30085106, 21701560, 33387674, 36212643, 29110115, 26825290, 27328748, 31868880, 33718295, 35914810, 28667181 |
| **Stalk A (middle):** G359A, G359R, G362D, G362S, G362D/I512T, G363D, F370C, A395D**, A395G, G401S, R403C, E410K, C431Y | epileptic encephalopathy, developmental delay, microcephaly, pain insensitivity, mild speech delay ataxia, muscle tone abnormalities, optic atrophy** | hyperfilamentous, elongated mitochondria, giant mitochondria, reduced mtDNA content, normal/abnormal peroxisome morphology | Verigni et al 2019, Trailo-Graovac et al 2019, Sheffer et al 2016, Vanstone et al 2016, Waterham et al 2007, Whitley et al 2018, von Spiczak et al 2017, Schmid et al 2019, Ladds et al 2018, Nolden et al 2022, Fahmer et al 2016, Vandelaar et al 2019, Ryan et al 2018 | 30801875, 31475481, 26992161, 26604000, 17460227, 30085106, 28667181, 30939602, 30109270, 35914810, 27145208, 31587467, 29877124 |
| **GED Domain:** Y691C, R710G, Q721Ter | epileptic encephalopathy, developmental delay, nystagmus, hypotonia | large, abnormal mitochondria and peroxisomes | Assia Batzir et al 2019, Nolden et al 2022, Zhang et al 2024 | 30850373, 35914810, 38341530 |

In this study, we report the results of using iPSC lines harboring reported dominant-negative heterozygous mutations in either the GTPase or stalk domain of DRP1 (G32A or R403C, respectively) (**Fig. 1A**) to generate cortical cultures and characterize the effects of disrupted mitochondrial fission on neuronal development and synaptic maturation. We found that iPSCs with DRP1 mutations generate cultures with a similar composition of cortical neurons compared to control. Mutant DRP1 neurons maintained highly elongated mitochondria within their axons throughout maturation, and we subsequently observed significant size-dependent changes to mitochondrial motility that were mutation specific. Cortical neurons were cultured for 35 and 65 days and then subjected to bulk RNA sequencing (RNA-seq). RNA-seq analyses demonstrated mutation-specific differences in synaptic and other developmental genes that matched strongly with observed patient and *in vitro* phenotypes, with new insights into possible underlying mechanisms of changes in neuronal activity. Mutant DRP1 neurons also had functional differences compared to control neurons, including deficits in synaptic development and more robust stimulated calcium responses.

**Figure 1.**
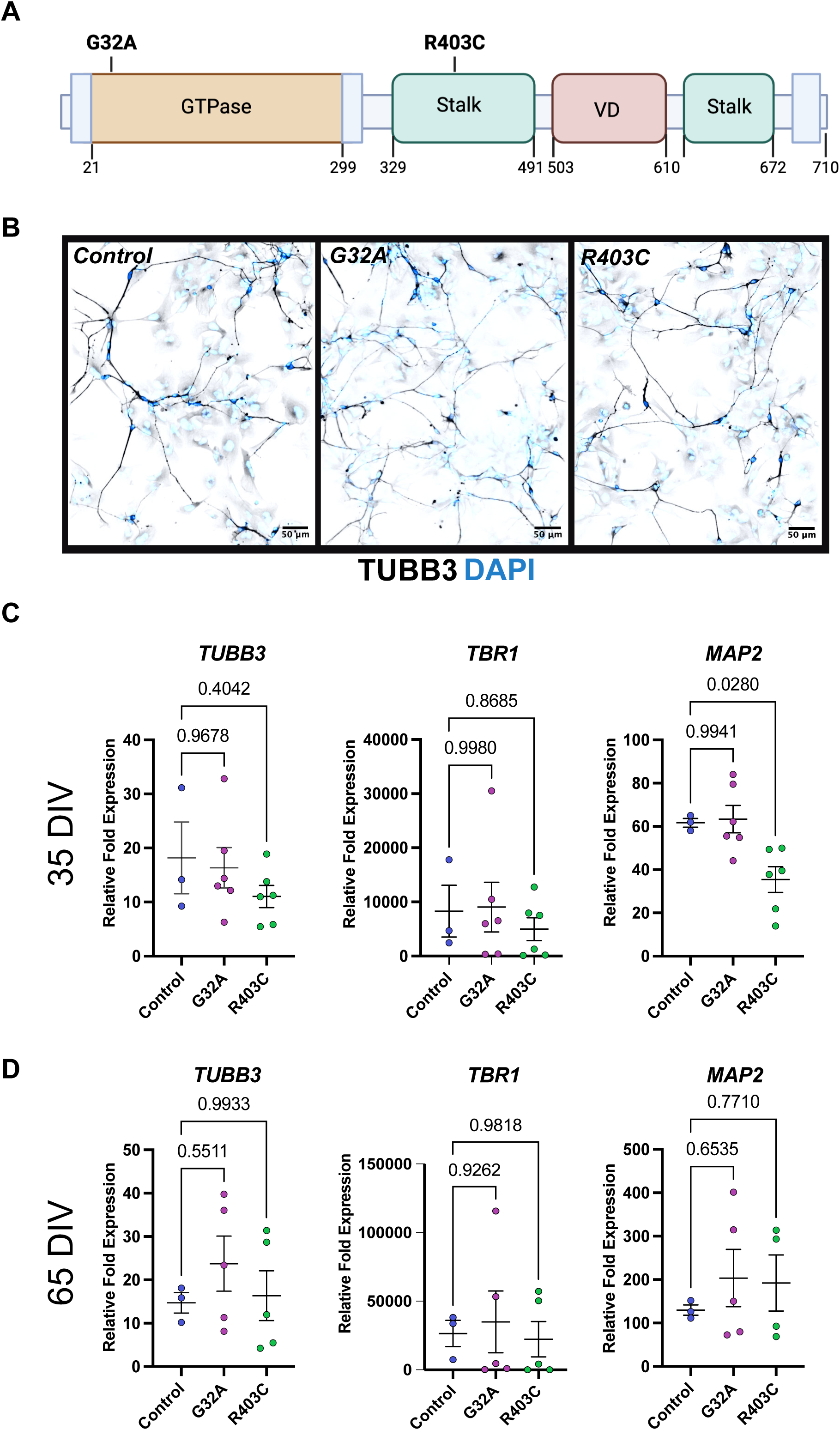
Characterization of mutant DRP1 cortical cultures. (**A**) Schematic of patient mutations in GTPase (G32A) or stalk (R403C) domain of DRP1 (**B**) Immunofluorescent staining of iPSC-derived 25 DIV cortical cultures for TUBB3/DAPI; scale, 50 µm (**C**) qPCR relative fold expression of markers of cortical differentiation TUBB3, TBR1, MAP2 in mutant DRP1 cultures compared to control at 35 and 65 DIV (one-way ANOVA; control n=3, mutant n=5; 35 DIV MAP2 F_2,10_ = 1.32, df=6.46, G32A p=0.99, R403C p=0.02; TUBB3 F_2,10_=0.43, df=1.06, G32A p=0.96, R403C p=0.4; TBR1 F_2,10_ = 0.20, df=0.35, G32A p=0.99, R403C p=0.86; 65 DIV MAP2 F_2,10_ = 1.33, df=0.36, G32A p=0.65, R403C p=0.77; TUBB3 F_2,10_=0.1.74, df=0.67, G32A p=0.55, R403C p=0.99; TBR1 F_2,10_ = 0.43, df=0.15, G32A p=0.92, R403C p=0.98). Error bars represent mean ± SEM.

## Results

### Differentiation of DRP1 mutant iPSCs into cortical neurons

DRP1 mutant iPSCs were differentiated into cortical neurons via small molecule dual-SMAD inhibition (**Fig. 1B; Supplementary Fig. S1**)^13,14^. Cortical neuron markers were quantified at 35 and 65 days in vitro (DIV) via qPCR (**Fig. 1C-D**). Transcriptional levels of *TUBB3* and *TBR1* were found not to be significantly different between mutant DRP1 and control cultures at 35 DIV, indicating similar levels of glutamatergic cortical neurons in each culture. *MAP2* levels were found to be lower in DRP1 R403C mutant cultures compared to control, suggesting a decrease in the number of neurons in these cultures (**Fig. 1C**). By 65 DIV, transcriptional levels of *TUJ1, TBR1, and MAP2* in all mutant DRP1 neuronal cultures were comparable to control (**Fig. 1D**).

### Hyper-elongation of axonal mitochondria in DRP1 mutant cortical neurons

Cells containing G32A or R403C mutations in DRP1 reveal a hyper-elongated mitochondrial phenotype resulting from an impaired capacity to undergo fission^15^. Because smaller, “fissed” mitochondria are a critical precursor to efficient mitochondrial trafficking within neurons, it was unclear if hyper-elongated mitochondria would have the capacity to be trafficked within axons. Control and mutant DRP1 cultures were fixed at 35 and 65 DIV and stained for TUBB3 and mitochondrial marker HSP60. Mitochondria contained within axons were imaged at super resolution and quantified for morphological differences between DRP1 mutant and control neurons. At 35 and 65 DIV, axonal mitochondria in mutant DRP1 neurons were found not to be significantly different in volume or surface area compared to control neurons (**Fig. 2A, B**); R403C (but not G32A) axonal mitochondria were significantly more elongated compared to control mitochondria (**Fig. 2B**).

**Figure 2.**
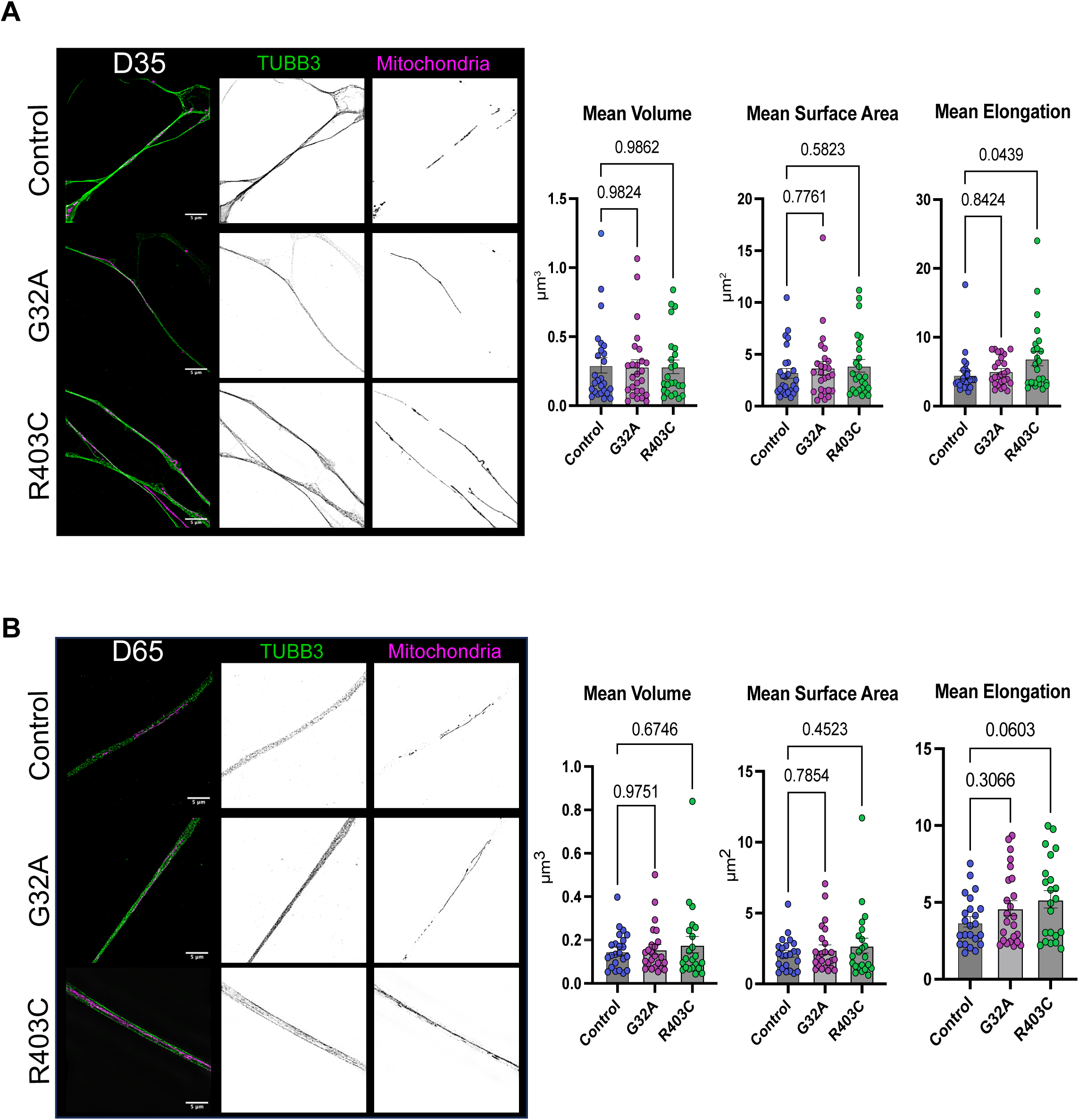
Morphology of mitochondria in axons. (**A**) SIM maximum intensity projection of immunofluorescent staining for mitochondria (anti-mito, magenta) in 35 and 65 DIV cortical axons (TUBB3, green). Right panels show inverted single channel images. Scale, 5 µm (**B**) Measurements of 3D axonal mitochondria morphology at 35 (left) and 65 (right) DIV averaged per axon (n = 24 axons, one-way ANOVA; 35 DIV volume F_2,73_ = 0.31, df=0.014, G32A p=0.98, R403C p=0.98; Surface Area F_2,73_ = 0.31, df=0.40, G32A p=0.77, R403C p=0.58; elongation F_2,73_ = 3.98, df=2.95, G32A p=0.84, R403C p=0.04; 65 DIV volume F_2,66_ = 0.88, df=0.30, G32A p=0.97, R403C p=0.67; Surface Area F_2,66_ = 0.1.21, df=0.59, G32A p=0.78, R403C p=0.45; elongation F_2,66_ = 3.18, df=2.41, G32A p=0.30, R403C p=0.06). Error bars represent mean ± SEM.

### Abnormal mitochondrial trafficking in DRP1 mutant cortical neurons

Live recordings were acquired at 65 DIV for control and mutant DRP1 cultures to visualize the dynamic movement and trafficking of DRP1 mutant mitochondria within axons (**Fig. 3**). Overall, mutant DRP1 neurons at 35 and 65 DIV display a significant increase in axons containing hyper-elongated mitochondria compared to control neurons (**Fig. 3A**). Axons from mutant DRP1 neurons were observed to have many dynamic mitochondria undergoing both anterograde and retrograde movement, stalling, and rapid directional change similar to control. Mitochondria within these axons were also observed to undergo some fusion and fission events (**Fig. 3A**, green arrow). Certain unique dynamic behaviors were observed frequently in mitochondria from mutant DRP1 neurons, such as extension and retraction of mitochondrial processes on hyper-elongated mitochondria without overall movement of the mitochondria itself (**Fig. 3A**, red arrow). Manual tracking of mitochondrial movement averaged across 10 axons per group showed no significant differences between mitochondria movement distance (average and total) or average velocity (**Fig. 3B**) in mutant DRP1 and control neurons. Axonal mitochondria from mutant DRP1 neurons were observed to be highly heterogeneous in size, with possible size-dependent differences in motility. Mitochondria were therefore binned by size (small, <2.5µm; medium, 2.5-7.5µm; large, 7.5-20µm; very large, 20+ µm) and analyzed again for differences in movement distance and velocity. Interestingly, small mitochondria within axons in R403C mutant neurons moved a significantly further average distance and with a faster average velocity compared to medium, large, or very large mitochondria in the same axons (**Fig. 3C**). Large mitochondria within axons in G32A mutant DRP1 neurons were found to move a significantly further total distance compared to small and medium sized mitochondria in the same axons. There were no significant differences in distance or velocity between mitochondria of different sizes in axons from control neurons (**Fig. 3C**).

**Figure 3.**
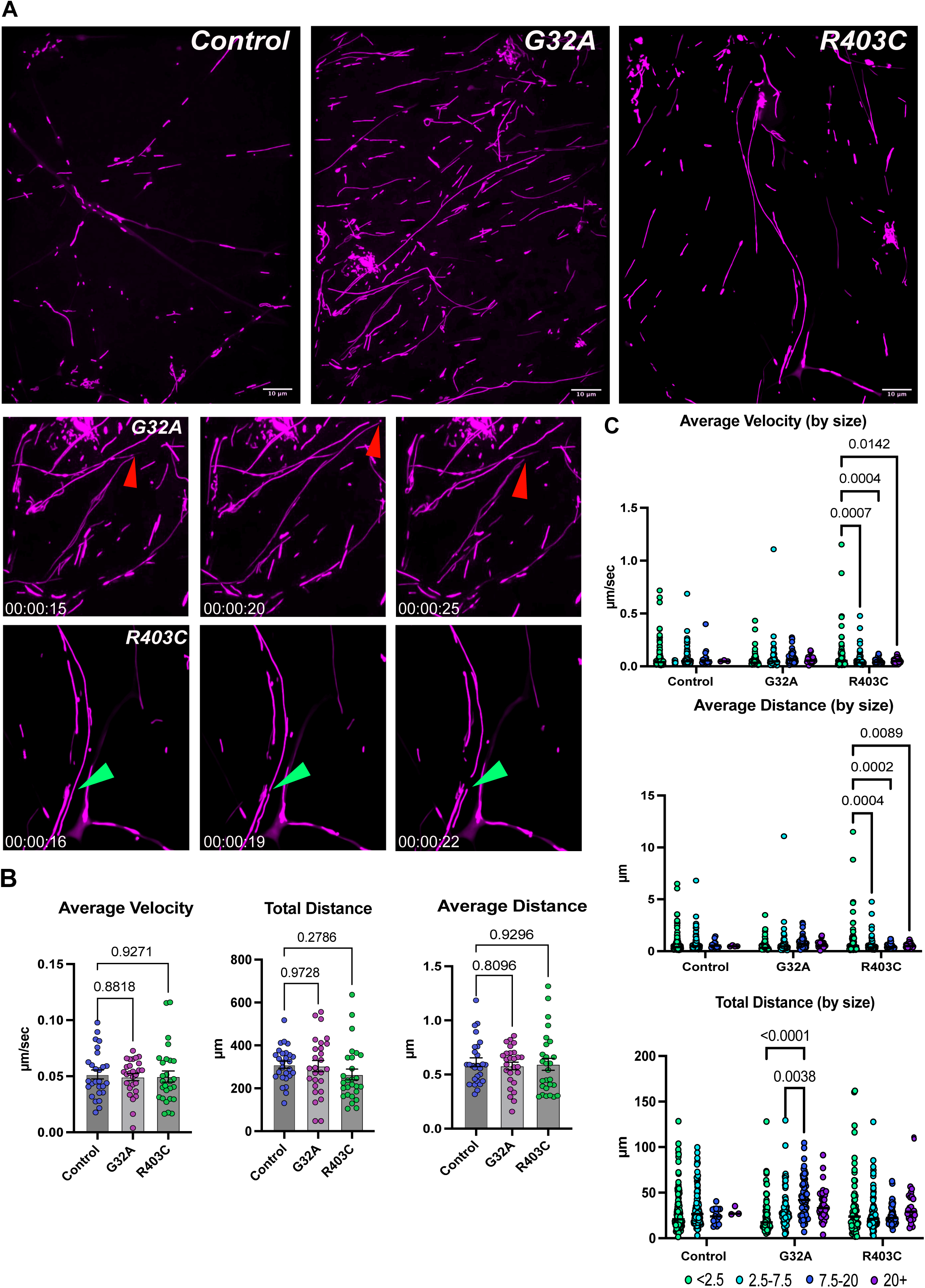
Motility of axonal mitochondria. (**A**) Spinning disk confocal max intensity projection of 65 DIV axonal mitochondria (magenta) recorded for 10 min show dynamics such as extension and retraction (red arrow) and fission events (green arrow); scale, 10 µm (**B**) Average distance, average velocity, and total distance of all axonal mitochondria movement averaged per axon (n=30 axons, one-way ANOVA; average distance F_2,78_=1.26, df=0.15, G32A p=0.80, R403C p=0.92;; total distance F_2,78_=1.85, df=1.16, G32A p=0.97, R403C p=0.27; average velocity F_2,78_=1.70, df=0.09, G32A p=0.88, R403C p=0.92). Error bars represent mean ± SEM (**C**) Average velocity, average distance, and total distance of axonal mitochondria movement binned by mitochondrial size (dots represent individual mitochondria, two-way ANOVA genotype x mitochondrial size, average distance F_6,736_=3.85, df=6, p=0.0009; total distance F_6,736_=5.137, df=6, p<0.0001; average velocity F_6,736_=3.85, df=6, p=0.0009; total distance F_6,736_=3.166, df=6, p=0.0045; p-values on graphs from Tukey’s multiple comparison post-hoc analysis). Bars represent median.

### RNA-seq analysis shows dysregulation of synaptic and ion channel genes in mutant DRP1 cultures

To better characterize differences in mutant DRP1 neuronal cultures during maturation, RNA was collected at 35 and 65 DIV and profiled with bulk RNA-seq analysis. Hierarchical clustering and principal component analysis (PCA) showed strong clustering between samples of the same timepoint (**Supplementary Fig. S2A-D**). PCA also showed strong clustering of both mutant DRP1 lines together, as well as strong separation from the control line (**Supplementary Fig. S2C**). These results indicate that mutations in different domains of DRP1 can result in overall similar changes in transcriptional landscape during neuronal development (**Supplementary Fig. S2**). Interestingly, both G32A and R403C 35 DIV mutant DRP1 lines showed more than 3000 differentially expressed genes (DEX) compared to control; however, by day 65, only ∼350 genes were found to be differentially expressed, indicating less transcriptional variability between mutant and control cultures over time (**Supplementary Fig. S2D**). At 35 DIV, the proteolipid neuronatin (*NNAT*), which has been implicated in ion channel regulation during brain development, was found to be the first and third most significant DEX gene in both R403C and G32A mutants compared to control, respectively (**Fig. 4A and 4B**)^16,17^. SLIT and NTRK Like Family Member 4 (*SLITRK4*), a member of the axonal growth controlling protein family is decreased at 35 DIV in both G32A and R403C cultures compared to control (**Fig. 4A**)^18^. RNA-seq results also reveal differences in gene expression between DRP1 mutant cultures, supporting a possible structure-function relationship between mutations in specific DRP1 domains and downstream patient phenotypes. Notable differences between DRP1 mutants include significantly decreased expression of protocadherin gamma C3 (*PCDHGC3*), a neural cadherin-like cell adhesion protein, and pyridoxamine 5’-phosphate oxidase (*PNPO*), an enzyme required for B6 (and downstream neurotransmitter) synthesis, in G32A cultures compared to R403C cultures at both 35 and 65 DIV (**Fig. 4A, and B**). Mutations in PNPO are known to be causal for an autosomal recessive pyroxidal-5’-phosphate (PLP) vitamin-responsive epileptic encephalopathy leading to refractory seizures during the first year of life^19,20^. Therefore, PNPO deficiency and some mitochondrial encephalomyopathy, lactic acidosis, and stroke-like episodes (MELAS) patients may have convergence of pathways related to disease etiology and possible therapeutic overlap. RNA-seq analysis also included comparing within mutant cultures over time between 35 and 65 DIV. Both DRP1 mutant cultures showed a significant increase in expression of complement C1q like 1 (*C1QL1*), which is associated with synaptic structure maintenance and remodeling, at day 65 compared to day 35 (**Supplementary Fig. 3**)^21^. This increase in *C1QL1* was not found in day 65 control cultures compared to day 35 cultures.

**Figure 4.**
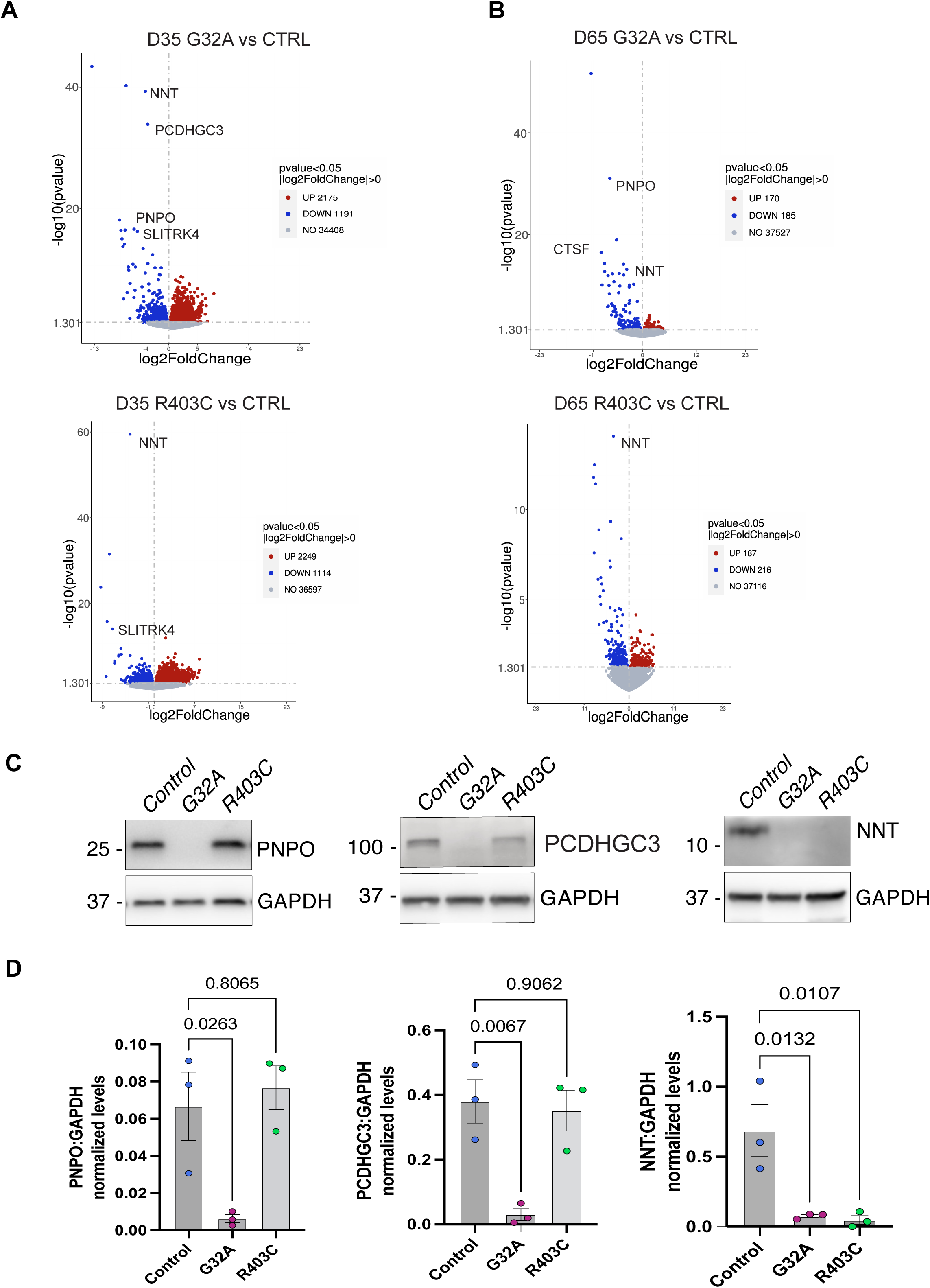
Bulk RNA-seq analysis of mutant DRP1 cultures. (**A**) Volcano plots of gene expression changes in 35 DIV G32A cultures compared to control (top panel) and 35 DIV R403C cultures compared to control (bottom panel) (**B**) Volcano plots of gene expression changes in 65 DIV G32A cultures compared to control (top panel) and 65 DIV R403C cultures compared to control (bottom panel) (**C**) Western blots of PNPO (left panel), PCDHGC3 (center panel), and NNT (right panel) in G32A and R403C cultures 100+ DIV (**D**) Quantification of western blots of PNPO (left panel), PCDHGC3 (center panel), and NNT (right panel) (n=3 blots; one-way ANOVA; PNPO F_2,6_=9.06, df=0.73, G32A p=0.026, R403C p=0.80; PCDHGC3 F_2,6_=12.99, df=0.55, G32A p=0.0067, R403C p=0.90; NNT F_2,6_=10.98, df=2.07, G32A p=0.01, R403C p=0.01). Error bars represent mean ± SEM.

Western blot was used to confirm if transcriptional changes of top DEX genes persist at the protein level in mutant DRP1 cultures at later timepoints (over 100 DIV). Protein levels of PNPO and PCDHGC3 were found to be significantly decreased in G32A mutant DRP1 cultures compared to control, with little to no protein levels detected (**Fig. 4C and 4D**). NNT was found to be significantly decreased in both G32A and R403C mutant DRP1 cultures compared to control (**Fig. 4C and D**). These findings recapitulate changes in gene expression detected via RNA-seq profiling at 35 and 65 DIV and confirm their persistence as mutant DRP1 cultures continue to mature past 100 days in culture.

### Mutant DRP1 neuronal DEGs correlate with reported patient phenotypes

Gene Ontology (GO) analysis was completed on RNA-seq data set (GO terms with padj < 0.05 are significant enrichment). Both 35 DIV mutant DRP1 cultures were highly associated with changes in expression of genes related to embryonic development, axon projection formation, and ion channel activity terms compared to control cultures **(Fig. 5A and 5B**); most enriched GO terms of 35 DIV cultures were correlated with increased gene expression (**Fig. 5A**). 35 DIV R403C mutant cultures had a notable decrease in expressed genes associated with the post-synapse and dendritic spines. By 65 DIV, R403C mutant cultures had additional top GO terms related to synaptic activity, particularly neurotransmitter binding, when compared to control **(Fig. 5C and 5D).** When transcripts of cultures were compared over time from 35 to 65 DIV, we found a striking difference between GO terms enriched in control compared to mutant DRP1 cultures. Control cultures were associated almost exclusively with terms related to decreases in DNA replication and cell cycle—typical of progressively postmitotic neuronal maturation **(Supplementary Fig. 4A).** This enrichment observed in control was not found in mutant DRP1 cultures, which shared a strong enrichment of gene expression changes and GO terms associated with synapse development, ion channel activity, and microtubule function (**Supplementary Fig. 4B and 4C; left panel**). Kyoto Encyclopedia of Genes and Genomes (KEGG) enrichment analysis showed a strong redundancy of calcium signaling and neuroactive ligand-receptor interaction pathways in all mutant DRP1 cultures compared to control and over time (**Supplement Fig. 4A, 4B, and 4C; right panel**).

**Figure 5.**
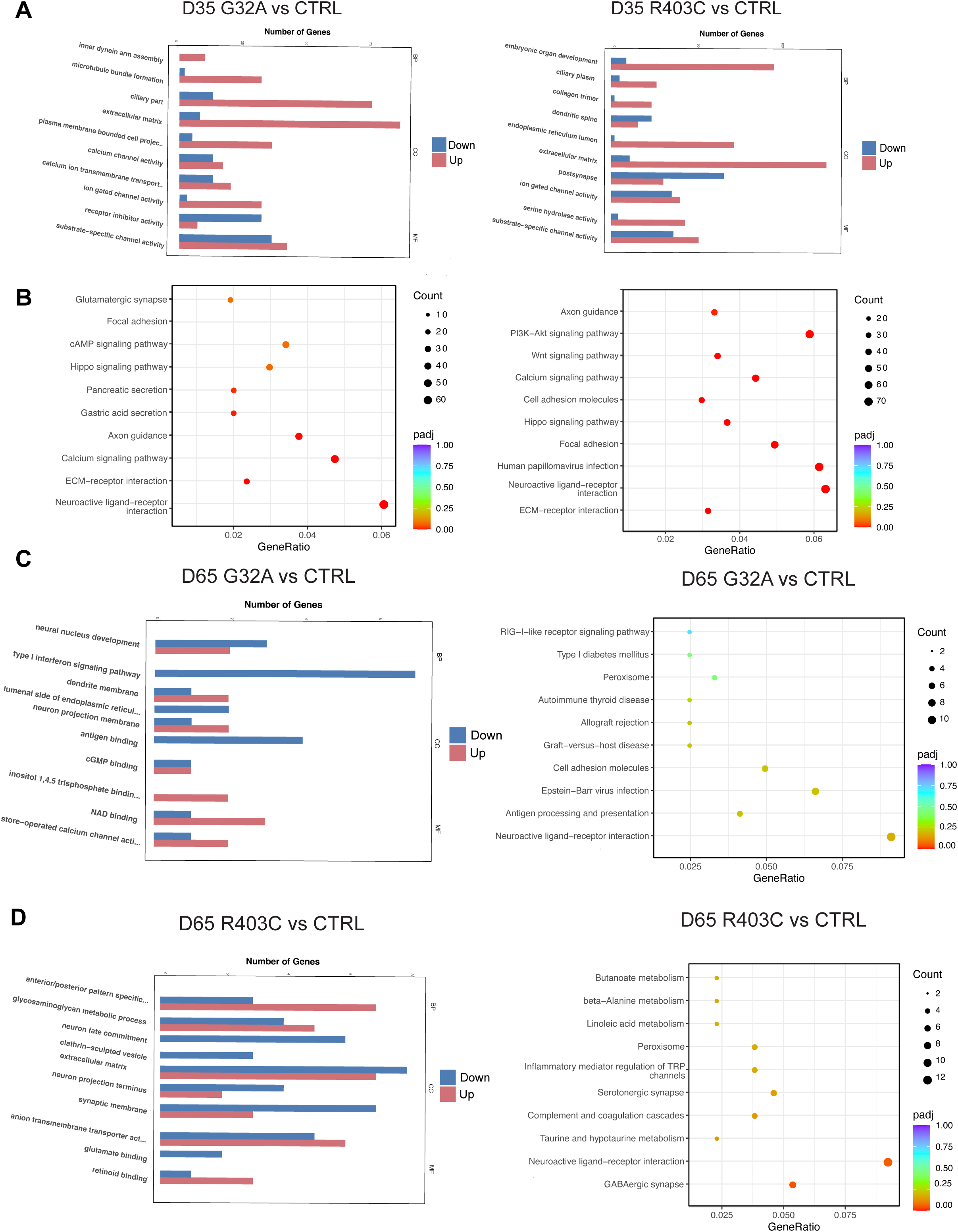
GO/KEGG enrichment analyses for mutant DRP1 cultures vs control. (**A**) top 10 GO terms enriched in 35 DIV G32A (left panel) and R403C (right panel) cultures (**B**) top 10 KEGG terms enriched in 35 DIV G32A (left panel) and R403C (right panel) cultures. (**C**) top 10 GO (left panel) and KEGG (right panel) terms enriched in 65 DIV G32A cultures (**D**) top 10 GO (left panel) and KEGG (right panel) terms enriched in 65 DIV R403C cultures.

### Disrupted synaptic development in DRP1 mutant cortical neurons

The results of RNA-seq analysis indicate a significant change in genes related to synaptic development and activity in DRP1 mutant cortical neurons. Indications of synaptic development changes are also supported by our evidence that axonal mitochondria in DRP1 mutant neurons have altered motility and are likely not being efficiently transported to synapses. Furthermore, presence of hyper-elongated mitochondria trafficking within axons suggests that the local mitochondrial network within synapses may not be able to meet the rapid changes to calcium and ATP demands as synapses develop and become active. During synaptic development pre- and post-synaptic structures develop proportionally to one another through activity-dependent positive feedback mechanisms^22,23,24^. To visualize differences in abundance and colocalization of pre- and post-synaptic proteins between mutant DRP1 and control neurons, cultures were fixed at 35 and 65 DIV and probed for pre-synaptic synaptophysin (SYP) and postsynaptic density protein 95 (PSD-95) markers using structured illumination microscopy (SIM) (**Fig. 6**). Visually, control DRP1 cultures appeared more synaptically dense with a more complex network of interconnected neurites (**Fig. 6A, B, C**). At both 35 and 65 DIV, pre-synaptic SYP volume was not significantly different; however, SYP of day 65 G32A mutant synapses were trending closely toward a significant decrease in volume (**Fig. 6B, C, E, F**). Post-synaptic PSD-95 volume was found to be significantly decreased at day 35 at G32A mutant synapses and significantly decreased by 65 DIV at R403C mutant synapses (**Fig. 6B, C, E, F**). Because pre- and post-synaptic structures develop concomitantly with each other, colocalization of SYP and PSD95 was quantified as a measure of overall synaptic strength^25^. Interestingly, a significant increase in overlap between SYP and PSD95 was found at G32A mutant synapses by day 35 compared to both control and R403C mutant synapses (**Fig. 6B, E**). By day 65, there were no significant changes in overlap of SYP and PSD95 at DRP1 mutant synapses compared to control (**Fig. 6C, F**). Overall, there was a notable increase in colocalization between SYP and PSD95 in all groups from day 35 to day 65 (∼10-15% to ∼25-30% mean overlap), indicating significant synaptic strengthening and network development over time in all cultures (**Fig. 6D**).

**Figure 6.**
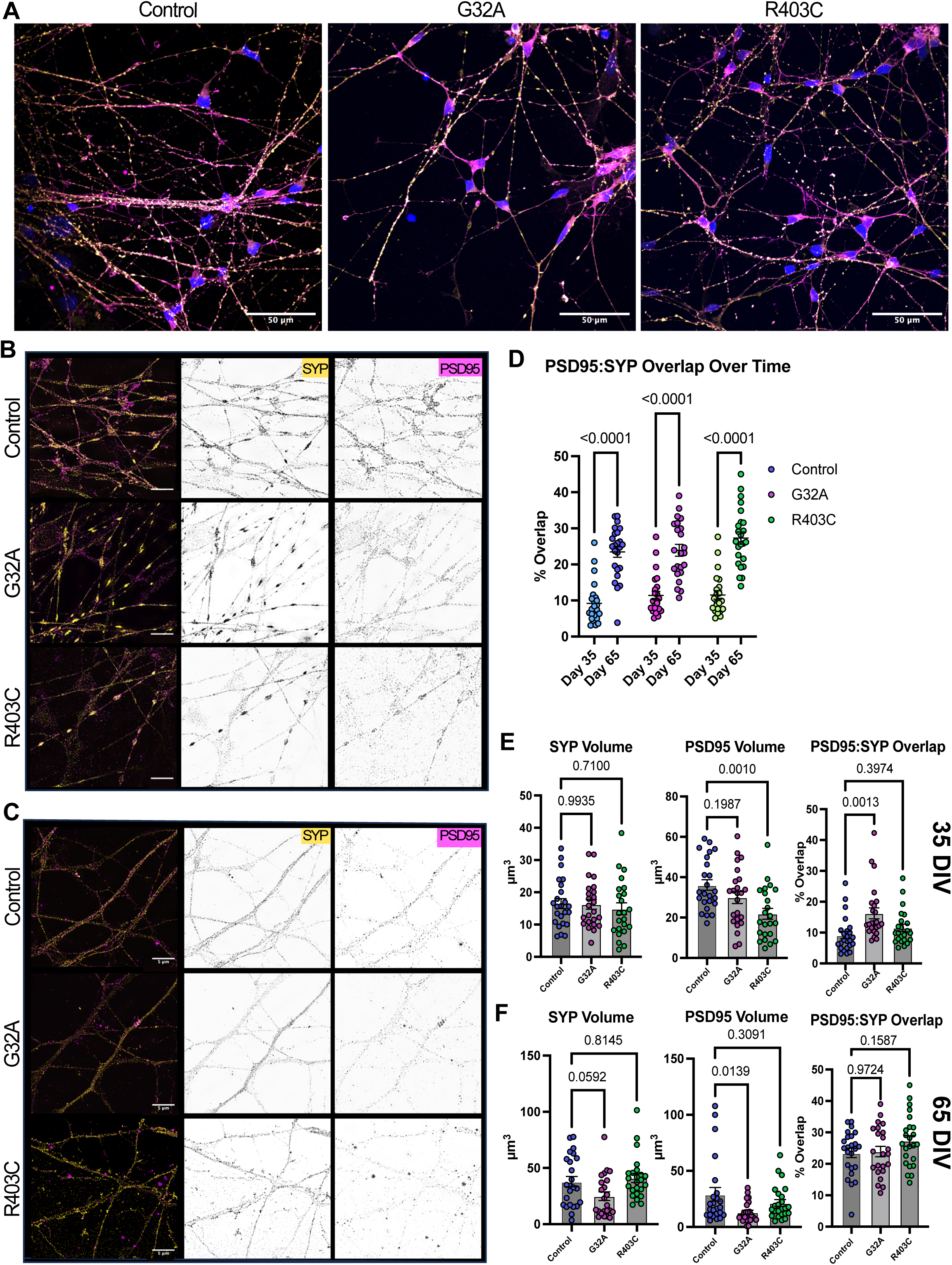
Synaptic development of mutant DRP1 neurons. (**A**) Spinning-disk confocal max intensity projections of immunofluorescent staining for SYP (yellow) and PSD95 (magenta) in 65 DIV cortical cultures; scale, 50 µm (**B**) SIM max intensity projections of immunofluorescent staining for SYP (staining) and PSD95 (magenta) in 35 DIV cortical cultures; scale, 5 µm (**C**) SIM max intensity projections of immunofluorescent staining for SYP (staining) and PSD95 (magenta) in 65 DIV cortical cultures; scale, 5 µm (**D**) Quantification of change in % overlap of SYP and PSD95 between 35 and 65 DIV (paired t-test, n=24 cells) (**E and F**) Quantification of synaptic density using SYP and PSD95 average volume and colocalization in 35 and 65 DIV cultures (n=24 cells; one-way ANOVA; 35 DIV SYP volume F_2,68_=0.41, df=0.28, G32A p=0.99, R403C p=0.71; PSD95 volume F_2,68_=0.17, df=6.6, G32A p=0.19, R403C p=0.001; colocalization F_2,68_=0.71, df=6.6, G32A p=0.001, R403C p=0.39; 65 DIV SYP volume F_2,68_=0., df=0., G32A p=0.059, R403C p=0.81; PSD95 volume F_2,68_=0., df=, G32A p=0.01, R403C p=0.3; colocalization F_2,68_=, df=, G32A p=0.97, R403C p=0.15). Error bars represent mean ± SEM.

### Altered calcium signaling activity of DRP1 mutant cortical neurons

Intracellular calcium, one of the most common second messengers in the brain, is tightly regulated in neurons as an important mediator of cell proliferation, death, and synaptic plasticity^26,27^. As action potentials arrive at the synapse, the activation of voltage-gated Ca2+ channels dictates the amount and duration of neurotransmitter release; mitochondria are essential regulators of intracellular Ca^2+^ ion compartmentalization, providing a buffer that maintains high sensitivity to Ca^2+^ fluctuations at synapses^28,29^. According to RNA-seq results, both G32A and R403C mutant DRP1 cultures show over a 2-fold decrease in expression of calcium voltage-gated channel auxiliary subunit gamma 3 (*CACNG3*) compared to control cultures at 65 DIV (data not shown). *CACNG3* is an AMPA receptor regulatory protein involved in modulation of trafficking and gating of AMPARs at the plasma membrane; *CACNG3* is reported to be important for slowing activation, deactivation, and desensitization of AMPARs.^30^ These results, as well as changes in pre- and post-synaptic markers, indicates a strong likelihood of perturbation of calcium dynamics in mutant DRP1 neurons. To visualize potential changes in calcium dynamics, 65 DIV cortical cultures were treated with cell permeant calcium dye Fluo-4 AM and imaged immediately using high resolution confocal microscopy. Cells were cultured in artificial cerebrospinal fluid (aCSF) media and recorded briefly to establish baseline activity, followed by subsequent treatment with 150 µM of glutamate (Glu) to stimulate neuronal activity (**Fig. 7**)^31^. Cultures were found to be spontaneously active at baseline without addition of agonist, and highly synchronously reactive post Glu addition (**Fig. 7A**). Mean fluorescence over time was calculated by averaging across 50 axonal ROIs both at baseline and after Glu stimulation per replicate (**Fig. 7B**). Mean amplitude of change in fluorescence after Glu stimulation was significantly higher in both G32A and R403C mutant groups, indicating a larger Ca^2+^ response compared to control (**Fig. 7C**). Mean time from stimulation to peak response trended lower for both mutant DRP1 cultures, with R403C approaching a significantly lower time compared to control (**Fig. 7C**). Area under the curve was significantly higher in G32A DRP1 cultures and approaching significantly higher in R403C DRP1 cultures compared to control (**Fig. 7C**). Overall, these findings indicate a more robust calcium response from mutant DRP1 cultures compared to control at 65 DIV.

**Figure 7.**
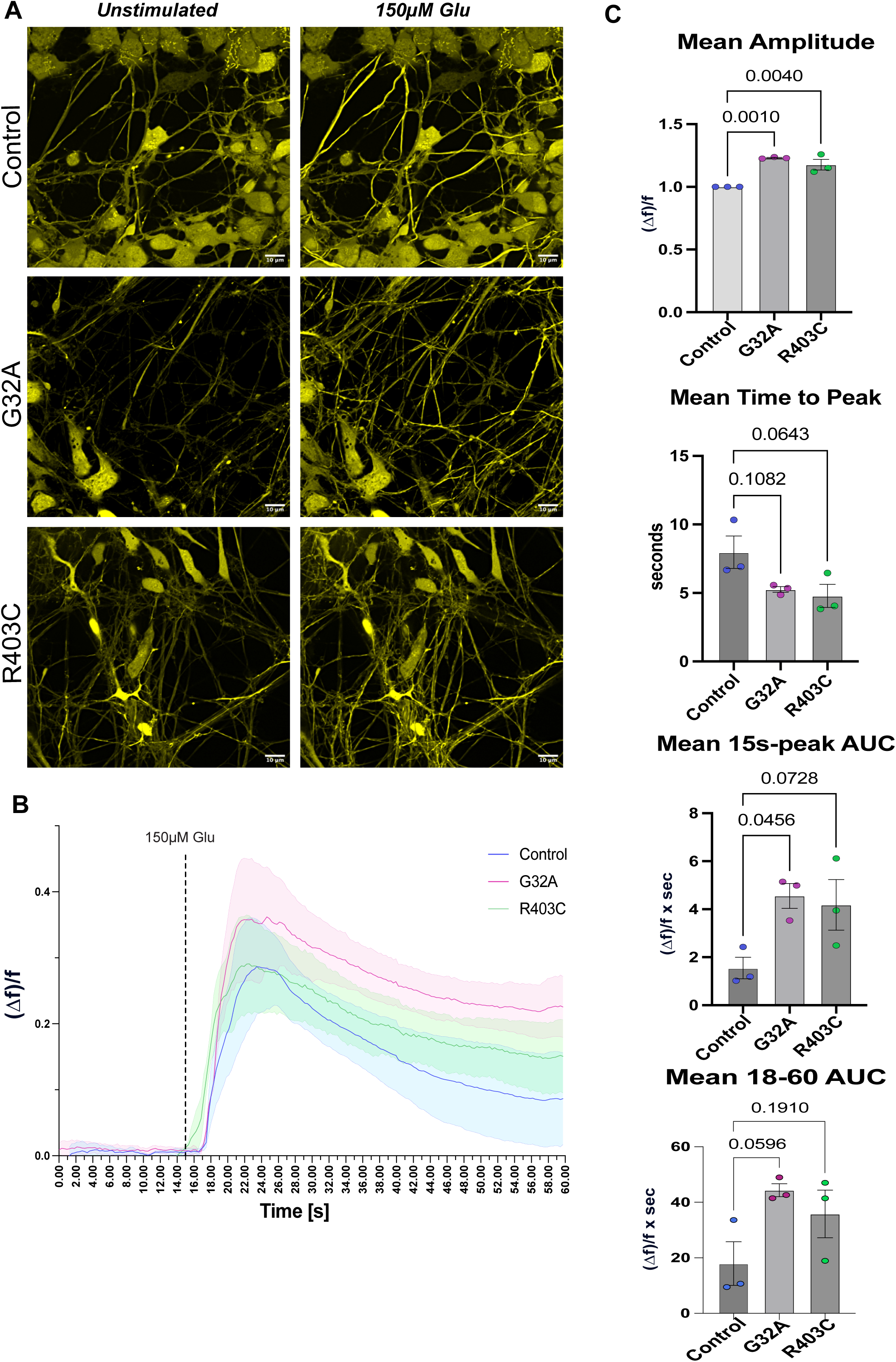
Calcium Response in Mutant DRP1 Cultures. (**A**) Calcium response (Fluo-4AM, yellow) in confocal single-plane images of 65 DIV cortical cultures recorded for 1 minute before (left) and after (right) 150 µM Glu stimulation in control and mutant DRP1 cells; scale, 10 µm (**B**) Change in fluorescence traces over time (average response across 50 ROIs per replicate) pre- and post Glu stimulation at 15s in control and mutant DRP1 cultures (C) Mean amplitude of change in fluorescence, mean time to peak after Glu stimulation, and area under the curve from Glu stimulation to peak response (individual ROIs normalized to control per replicate, one-way ANOVA; mean amplitude F_2,6_ = 1.85, df=24.23, G32A p=0.001, R403C p=0.004; mean time to peak F_2,6_ = 0.41, df=4.14, G32A p=0.10, R403C p=0.06; mean 15s to peak AUC F_2,6_ = 0.65, df=5.12, G32A p=0.04, R403C p=0.07; mean 18-60 AUC F_2,6_ = 3.87, df=0.38, G32A p=0.059, R403C p=0.19) of control and mutant DRP1 calcium traces. Error bars represent mean ± SEM.

## Discussion

It is well established that highly dynamic mitochondria are essential for proper neurodevelopment, with multiple studies examining the complex quality control mechanisms that ensure the proper distribution and movement of mitochondria within axons^32,33,34^. It is therefore of interest to understand how neurons with fission defects can survive during early stages of neurodevelopment. High and super-resolution imaging of neurons with mutant DRP1 reveals the capacity for highly elongated mitochondria to be actively trafficked within axons and to exhibit some dynamic behaviors, such as elongation and fragmentation of the network. Rapid directional movement, both anterograde and retrograde, indicates that these elongated mitochondria are actively trafficked, likely via the canonical axonal motor proteins kinesin and dynein on microtubule highways. Elongated mutant DRP1 mitochondria were also observed to extend and retract in a stretching motion, without lateral movement of the whole mitochondria. This motility behavior could be linked to an attempt and failure of microtubule motors to transport some very large mitochondria cargo. Studies have shown that neurons have quality control mechanisms which help regulate the size of mitochondria traveling within axons^35,4^. As it is currently unclear at what size mitochondria are incapable of being transported, these fission-defective cells provide a new opportunity to study the transport of large cargo within axons. It is possible that disruption of proper mitochondrial motility interferes with effective transport of other cargoes within the axon, further hindering necessary cell components from reaching the synapse. Mutant DRP1 axonal mitochondria were found to be highly variable in size with significantly increased numbers of large (7.5 – 20 µm) and very large mitochondria (20+ µm) compared to control axons; R403C mitochondria were significantly more elongated compared to control mitochondria. Live recordings indicated changes in mitochondria motility are size-dependent, with small R403C mitochondria moving further and faster on average than any other larger mitochondria within the same axons. Additionally, large G32A mitochondria moved further overall compared to smaller mitochondria within the same axons. These mutation-specific differences in mitochondrial motility (which are absent in control axons) indicate a possible compensatory response that allows mutant DRP1 neurons to leverage their highly variable mitochondrial size caused by fission disruption to increase mitochondrial transport and ultimately cell survival.

RNA-seq results of 35 and 65 DIV cultures demonstrate notable changes in transcriptional profiles when comparing mutant DRP1 cultures to control. Our hypothesis that cortical development and synaptic maturation are affected in a mutation-specific manner is further supported by notable loss in expression of PNPO and PCDHGC3 in G32A, but not R403C cultures. PCDHGC3 isoforms have been found to be required for control of synapse formation, size, and peripheral branching in developing mouse dorsal root ganglia^36^. Additionally, multiple studies have linked gamma protocadherin function to synaptic maturation and stabilization; mutations in these genes, such as protocadherin-19 (PCDH19), have to found to increase to seizure susceptibility^37,38,39^. Our findings provide further evidence of gamma protocadherins as possible risk genes for developing childhood encephalopathies.

The PNPO enzyme catalyzes the terminal, rate-limiting step in the synthesis of pyridoxal 5’-phosphate, or vitamin B6, a critical cofactor for neurotransmitter synthesis. It is currently unclear how disruption of the DRP1 GTPase domain activity connects mechanistically to vitamin B6 synthesis. Mutations in *PNPO* have been linked to a similar form of neonatal epileptic encephalopathy, PNPO deficiency^20^, in which patients respond to pharmacologic supplementation with pyridoxal 5’-phosphate (PLP) and/or pyridoxine^40,41,42^. Vitamin B6 supplementation could provide a potentially low-risk therapeutic to patients harboring mutations in specific domains of DRP1. Changes in PNPO and PCDHGC3 at the transcript and protein level provide evidence that defects in specific domains of DRP1 lead to mutation-specific cellular dysfunction that may result in variable patient phenotypes and therapeutic outcomes.

Changes in gene expression of *NNT*, however, were found in both G32A and R403C cultures compared to control, suggesting a convergent mechanism of overall DRP1 dysfunction that leads to changes in gene expression. NNT is a proteolipid that has been implicated in regulation of ion channels during brain development^43,44^. Though its specific role remains unclear, NNT appears to help regulate intracellular calcium homeostasis. NNT has been associated with Lafora disease (LD) a teenage-onset inherited progressive myoclonus epilepsy characterized by accumulations of intracellular glycogen inclusions called Lafora bodies^17,45^. Decreased expression of NNT in both DRP1 mutants may contribute to a more robust, unconstrained calcium response in neurons leading to hyperexcitability of synapses and epileptic phenotypes^17,43^.

Synaptic maturation was assessed in mutant DRP1 cultures using super-resolution microscopy of synaptic structures and measurements of functional activity via calcium recording. At 35 DIV, a significant increase in percent overlap between pre- and post-synaptic markers of G32A compared to control synapses indicates a higher density of synapses in these mutant cultures. Ultrastructural studies of synapse development indicate there are mechanisms that ensure as pre- and post-synapses are modified, they remain proportional to each other^23,22^. Changes in postsynaptic volumes can subsequently be interpreted as a response to concentrations of neurotransmitter release from the pre-synapse^46,47^. Decreases in average volume of G32A 65 DIV synapses (SYP and PSD95 markers) could therefore indicate a possible decline in synaptic activity and overall weakening of these synaptic connections. By 65 DIV, G32A cultures had normalized to a similar synaptic density compared to control. R403C post-synapses were found to be significantly decreased in volume at 35 DIV; however, this change was not persistent up to 65 DIV where post-synaptic volume was similar to control. Patterns of typical synaptic development have shown that weak synapses are less likely to survive, resulting in a positive selection for stronger synapses^48,49^. It is likely that cells with weak post-synaptic growth at earlier timepoints are less likely to survive in R403C cultures, with only more proportionately developing synapses remaining by 65 DIV.

Calcium recordings of 65 DIV mutant DRP1 cultures showed a significantly increased peak amplitude of intracellular calcium transients in response to glutamate stimulation. In addition, both mutant groups showed decreased time to peak response from stimulation, with R403C cultures having a significantly faster response time compared to control. This increase in rate and magnitude of calcium response may be related to significant downregulation of *CACNG3*, which aids in inhibitory control of AMPARs. It is notable that *CACNG3* is a reported risk gene for a form of childhood epilepsy^50^. Without this inhibition it is possible that dysregulation of channel opening leads to faster and increased influx of calcium into the cells, resulting in a more robust fluorescence response in DRP1 mutant cells compared to control. Calcium influx into the cell is tightly controlled to maintain intracellular sensitivity to acute fluctuations in calcium concentration necessary for neuronal activity. The mitochondrial network in G32A and R403C mutant DRP1 neurons is fission defective, and therefore, much less dynamic than a typical neuronal mitochondrial network. Dynamic mitochondria are required for effective calcium buffering in the cell, so mutant DRP1 neurons are likely less capable of compensating for intracellular calcium dysregulation. This is supported by our findings that mutant DRP1 cells show an increased area under the curve from peak to 60 seconds (G32A cells having a significantly higher area), which indicates a slower clearance of calcium from the cytoplasm. While calcium efflux is mainly mediated via transporter and channel activity, including the endoplasmic reticulum, calcium buffering via mitochondria is also important in this process.

In summary, this study used iPSC-derived cortical cultures with CRISPR induced heterozygous patient mutations in either the GTPase or stalk domain of DRP1 to study neuronal maturation and synaptic function. Our main findings show significant perturbations to mitochondrial trafficking, synaptic density, and calcium activity during differentiation and development of these cultures occurring in a mutation-specific manner. Notable changes to the transcriptional landscape of mutant DRP1 cultures identify risk genes commonly associated with other forms of childhood encephalopathies, highlighting converging cellular pathways leading to similar synaptic disorders. Growing understanding of these susceptibility loci increases the likelihood of early detection and access to appropriate therapeutic treatments for affected patients. Identifying DRP1 mutation-specific cellular phenotypes may help contribute to more personalized therapeutics to treat specific symptoms and increase lifespan of adolescents with these and other forms of encephalopathies.

## Methods

### Key Resources Table

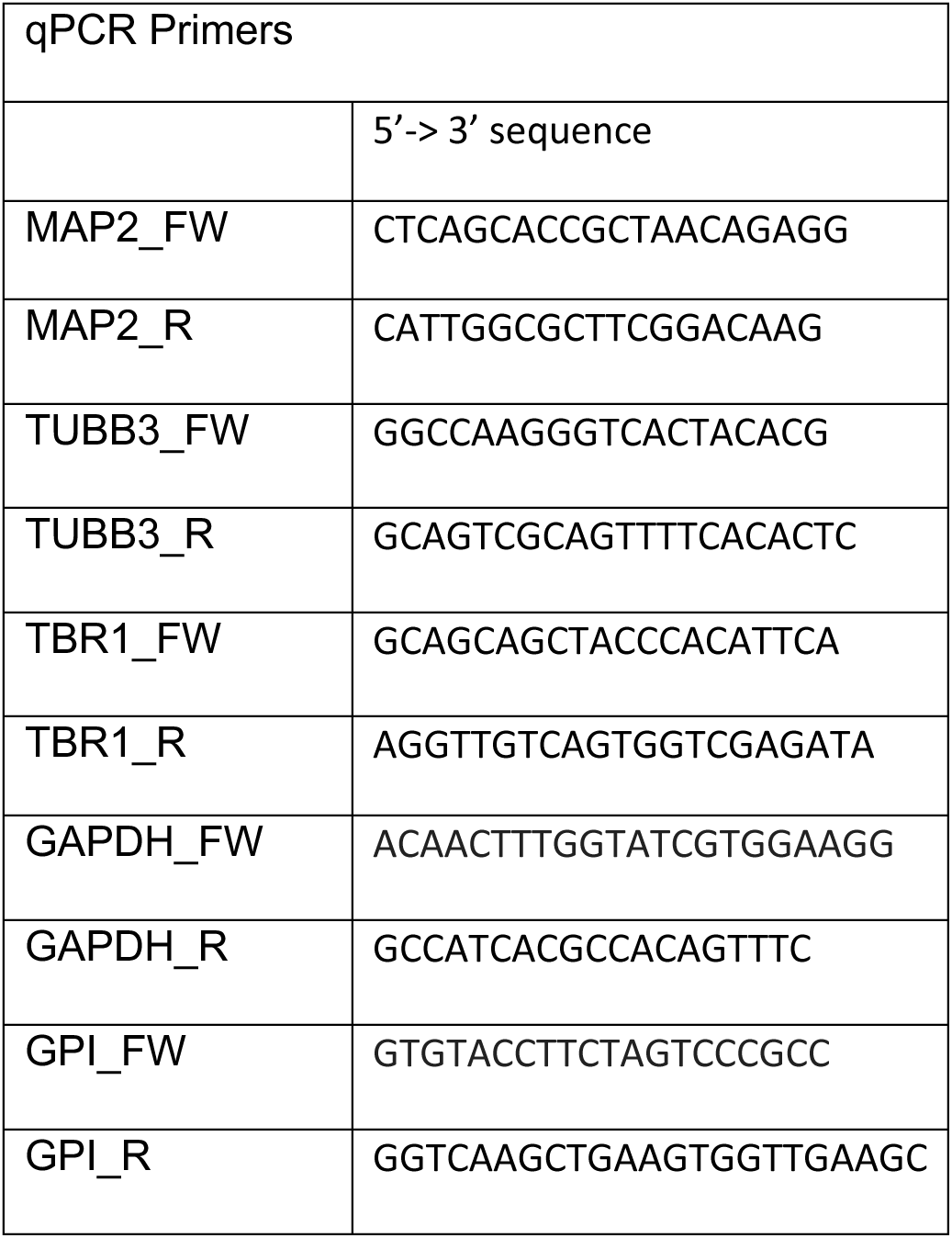

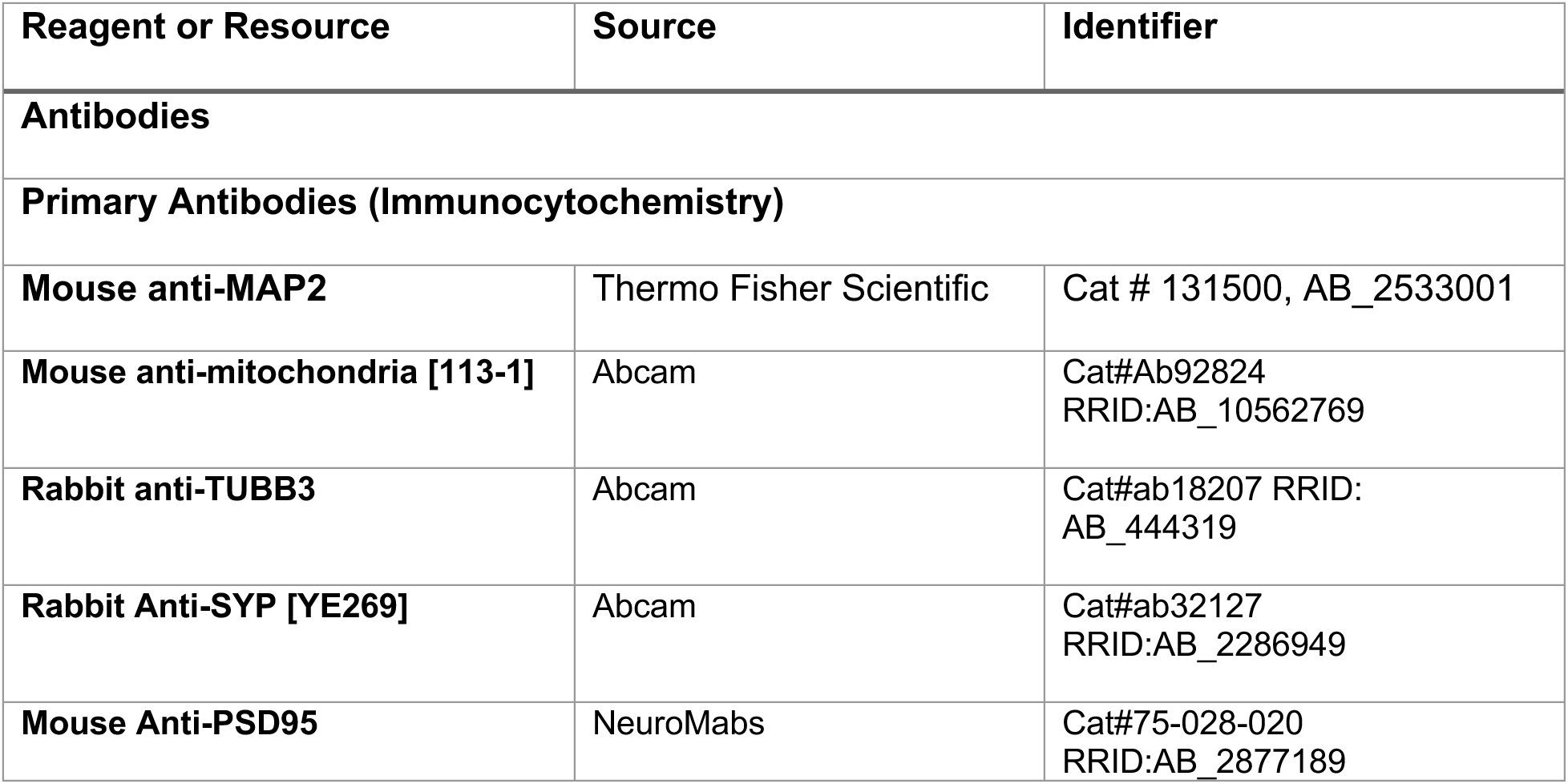
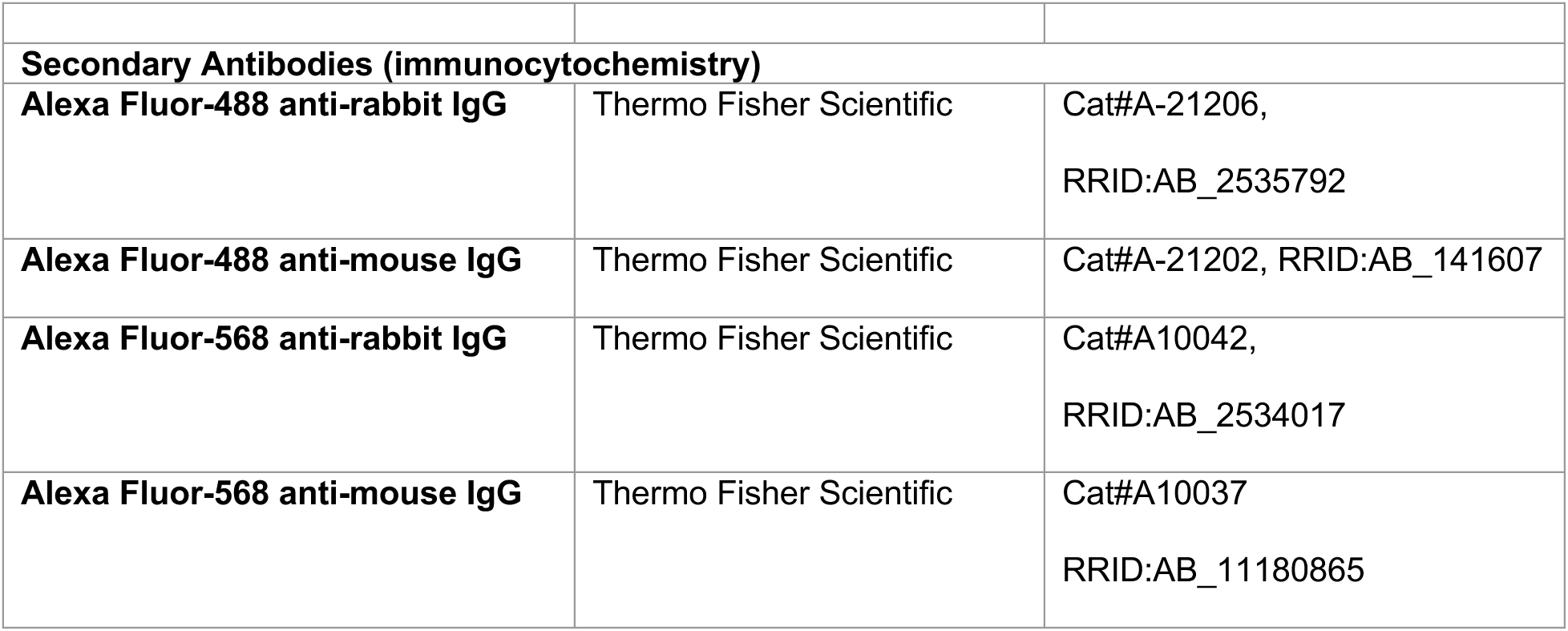

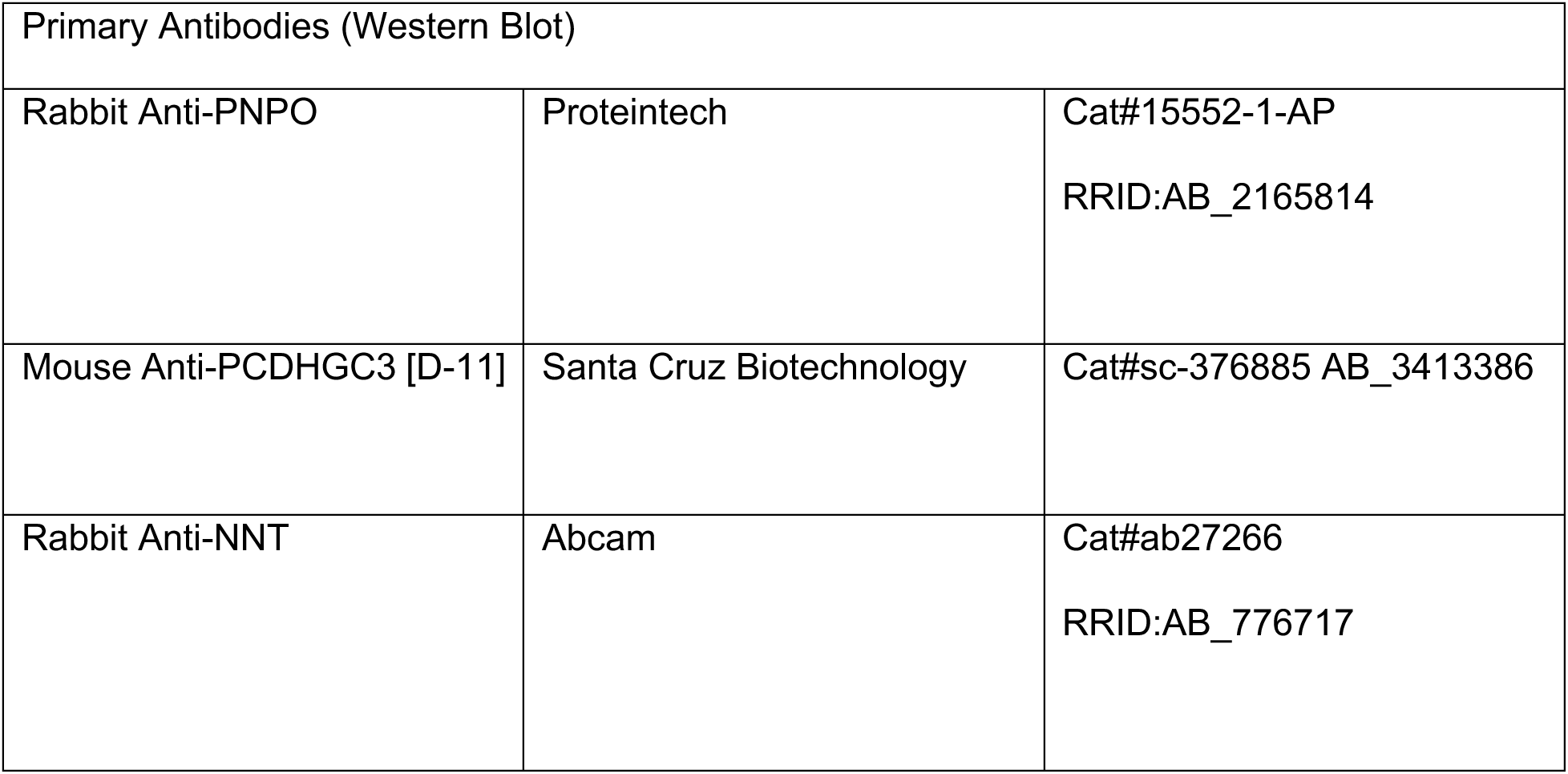

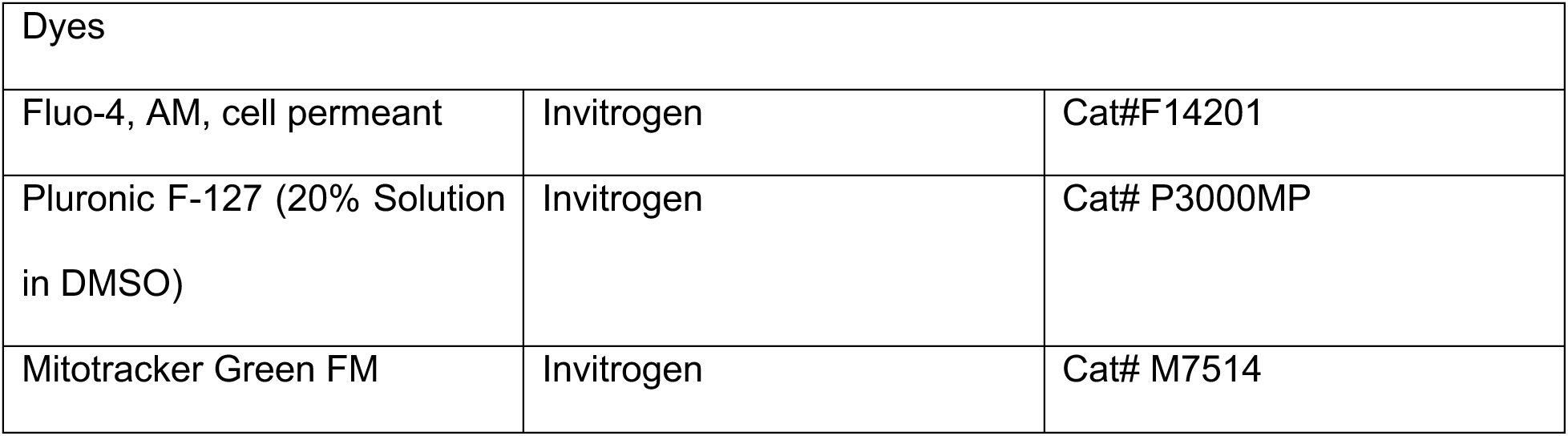

### Cell lines

The PGP-1 (GM2338) iPSC line used in this study was edited for DRP1 studies. The GM23338 (RRID: CVCL_F182) line is from a 55-year-old healthy, Caucasian male. In addition to one control cell line, two clones were used for each mutation (G32A and R403C). The cells were maintained on Matrigel-coated plates (Corning #354277) and cultured in StemFlex media (Thermo Fisher Scientific #A3349401) at 37°C with 5% CO2. Culture medium was changed daily. Cells were checked daily for differentiation and were passaged every 3-4 days. For passaging, iPSCs were dissociated with Gentle Cell Dissociation (Stem Cell Technologies #100-0485) and incubated at room temperature for 4 minutes. All experiments were performed under the supervision of the Vanderbilt Institutional Human Pluripotent Cell Research Oversight (VIHPCRO) Committee. Cells were checked for contamination periodically.

### Differentiation of iPSCs into cortical neurons

Cortical neurons were generated from PGP-1 iPSCs using the dual SMAD neural induction protocol published by Chambers et al. (2009; PMID: 19252484), using 10mM of SB4 and modified to use 0.4μM LDN (Stemgent #04-0074) instead of Noggin. The iPSCs were dissociated via incubation in Accutase (Innovative Cell Technologies #AT104) for 8 minutes at 37°C and 5% CO_2_. They were then resuspended in StemFlex (Thermo Fisher Scientific #A3349401) media with 10μM of ROCK inhibitor (Y27632; Stem Cell Technologies #72304) and centrifuged at 200x*g* for 3 minutes. Cells were plated at 100,000 cells/mL, and neural induction was started when iPSCs reached 100% confluency. For days 0-4 of differentiation, cells are maintained in neuralization media: 410mL Knockout DMEM/F12 (Invitrogen #12660012), 75mL Knockout Serum Replacement (Thermo Fisher Scientific #108280238), 5mL Glutamax (Thermo Fisher Scientific #35050061), 5mL Penicillin-Streptomycin (Thermo Fisher Scientific #15070063), 5mL MEM Non-Essential Amino Acids (Thermo Fisher Scientific #11140050), and 1.93μL of β-mercaptoethanol (BioRad #1610710).

Beginning on day 5, cells are maintained in an increasing concentration of N2 media (Supplementary Fig. 1) 500mL DMEM/F12 (Thermo Fisher Scientific #11330032), 0.775g D-Glucose (Sigma Aldrich #, and 5mL N2 Supplement (Thermo Fisher Scientific #17502048).

Neural differentiation is continued after 10 days of neural induction (Shi et al). After day 10, cells are maintained in 50% N2 CTX media and 50% B27 Neurobasal media. N2 CTX media: 500mL DMEM/F12 + Glutamax (Thermo Fisher Scientific #10565018), 5mL N2 supplement, 5mL MEM Non-Essential Amino Acids, and 1.93μL of β-mercaptoethanol. B27 neurobasal media: 500mL Neurobasal Media (Thermo Fisher Scientific #21103049), 10mL B27 supplement (Thermo Fisher Scientific #), and 5mL Glutamax (Thermo Fisher Scientific #17504044).

Cells were replated on days 21 and 35 at 5of differentiation at 500,000 cells/mL for downstream experiments.

At day 21, cells were replated at 5e5 cells/ml; cells planned for calcium assays were co-cultured with P1 mouse glia at an ∼ 3:1 ratio (neurons:glia).

### Immunofluorescence

Day 35 cells were fixed with 4% paraformaldehyde for 20 min and permeabilized in 1% Triton-X-100 for 10 min at room temperature. After blocking in 10% BSA, cells were treated with primary and secondary antibodies using standard methods. Day 65 cells were fixed in 4% paraformaldehyde for 30 min at 4C and permeabilized/blocked in 5% Donkey Serum, 0.3% Triton X-100 for 1 hour at room temperature. Cells were mounted in Fluoromount-G (Electron Microscopy Sciences #17984-25) prior to imaging. Primary antibodies used include mouse anti-Mitochondria (Abcam #ab92824), anti-TUBB3 (Abcam #ab18207), anti-SYP (Abcam #ab32127), anti-PSD95 (Neuromabs #75-028-020), anti-MAP2 (Thermo #131500). Secondary antibodies used include donkey anti-mouse Alexa Fluor-488 (Thermo Fisher Scientific #A21202), donkey anti-rabbit Alexa Fluor-488 (Thermo Fisher Scientific #A21206), and donkey anti-mouse Alexa Fluor-568 (Thermo Fisher Scientific #A10036), and donkey anti-rabbit Alexa Fluor-568 (Thermo Fisher Scientific #A10040). Super-resolution images were acquired using Nikon SIM microscope equipped with a 1.49 NA 100x Oil objective and Andor DU-897 EMCCD camera. Quantification of microscopy images was performed in NIS-Elements (Nikon) or Fiji. For mitochondrial morphology an axonal ROI was manually drawn followed by segmenting mitochondria in 3D and performing skeletonization of the resulting 3D mask. Skeleton volume, surface area, and elongation measurements were exported into Excel.

### Immunoblots

Cells were harvested in lysis buffer: 1% Triton, 10X phosphatase inhibitor (PhosSTOP; Roche, Sigma Aldrich #4906837001), 10x protease inhibitor (PIC; roche, Sigma Aldrich #04693159001), and 100X phenylmethylsulfonyl fluoride (PMSF) (RPI #P20270). Lysate protein concentrations were quantified using BCA analysis (Fisher Scientific #23225). Western blotting was performed using the Bio-Rad Mini-PROTEAN Electrophoresis System (Bio-Rad #1658036). Samples were run on either 4-20% or 12% Mini-PROTEAN TGX gels (Bio-Rad #4561094/#4561044). Protein was transferred from gels onto methanol-activated PVDF membranes (Bio-Rad #1620177). All membranes were washed in Tris-buffered saline with 0.1% Tween-20 (TBST) and blocked in TBST containing 5% milk. Primary antibodies were incubated at 1:1,000 overnight at 4°C unless otherwise stated: [List antibodies]. The following secondary antibodies were used at 1:10,000 dilution and incubated for 1 hour at room temperature in 5% milk w/ TBST: HRP donkey anti-rabbit (Jackson ImmunoResearch #711-035-152) and HRP goat anti-mouse IgG subclass 1-specific (Jackson ImmunoResearch #115-035-205). Protein signal was developed using either Pierce ECL Western Blot Substrate (Thermo Fisher Scientific #32109) or SuperSignal West Femto (Thermo Fisher Scientific #34094), depending on the sensitivity of the primary antibody. Protein signal was visualized using the Amersham Imager 600 (GE Life Sciences #29-0834-61). The obtained images were quantified using Fiji (imagej.net/software/fiji/), and representative images were composed using Affinity Designer 2 (affinity.serif.com/en-us/designer/).

### RNA extraction and cDNA synthesis

Cells were washed with PBS and scraped into 1000μl Trizol reagent. 200μl of chloroform was added, and the samples were incubated at room temperature for 5 minutes. The samples were centrifuged at 12,000 g, and the aqueous phase was collected. 500μl of isopropanol was added to precipitate RNA, and the sample was incubated for 25 minutes at room temperature, followed by centrifugation at 12,000 g for 10 minutes at 4°C. The RNA pellet was washed with 75% ethanol, semi-dried, and resuspended in 30μl of DEPC-treated water. After quantification and adjustment of the volume of all the samples to 1μg/μl, the samples were treated with DNAse (New England Biolabs #M0303). 10μl of this volume was used to generate cDNA using the manufacturer’s protocol (Thermo Fisher #4368814).

### Quantitative RT PCR (RT-qPCR)

1ug of cDNA sample was used to run RT-qPCR for the primers mentioned in the key resources table. QuantStudio 3 Real-Time PCR machine, SYBR green master mix (Thermo Fisher #4364346), and manufacturer instructions were used to set up the assay.

### Bulk RNA-sequencing

Raw reads of fastq format were firstly processed through Novogene in-house perl scripts. Clean reads were obtained by removing reads containing adapter, poly-N and low-quality reads from raw data; Q20, Q30 and GC content of the clean data were calculated. All downstream analyses were based on the clean data with high quality. Reference genome and gene model annotation files were downloaded from genome website directly. Index of the reference genome was built using Hisat2 v2.0.5 and paired-end clean reads were aligned to the reference genome using Hisat2 v2.0.5. featureCounts [3] v1.5.0-p3 was used to count the reads mapped to each gene; FPKM of each gene was calculated based on the length of the gene and reads count mapped to this gene. Differential expression [5] analysis was performed using DESeq2 (1.20.0). Adjusted p-values were calculated using the Benjamini-Hochberg False Discovery Rate (FDR) method. Genes with an adjusted P-value<=0.05 found by DESeq2 were assigned as differentially expressed. Gene Ontology [7] (GO) enrichment analysis of differentially expressed genes was implemented using clusterProfiler. GO terms with corrected P-value less than 0.05 were considered significantly enriched by differential expressed genes. clusterProfiler was used to test the statistical enrichment of differential expression genes in KEGG[8] pathways.

### Calcium Imaging

Fluo-4AM loading protocol adapted from Thermofisher’s protocol for fluo-4-based measurements of cytosolic calcium changes in neural stem cells in response to neurotransmitter application. Day 65 neuronal cultures co-cultured with P1 mouse glia were incubated with 2.86µM Fluo-4-AM cell permeant dye (Invitrogen F14201) with Pluronic F-127 solution (Invitrogen P10020) in aCSF for 1 hour at 37C. Cells were washed once with media followed by fresh aCSF media and imaged immediately. Cells were imaged on the Nikon W1 Spinning Disk Confocal 100X Plan Apo 1.49 NA Oil objective and a Prime 95B CMOS camera for 1 minute with 150µM Glutamate (Glu) (Thermo A15031) with stimulation occurring at 15s. Four wells were imaged per line in each replicate. Live recordings were processed with Richardson-Lucy deconvolution (20 iterations) and denoised using Nikon Elements software. Approximately 50 regions of interest (ROIs) were chosen across all recordings per group; Elements “Tracking” function was used to track mean fluorescence over time across all binaries. Change in fluorescence values were calculated as a function of change in fluorescence over baseline fluorescence (f/f) plotted over time. For statistical analysis, each G32A and R403C trace was normalized to a matched control for each replicate.

### Statistical Analysis

All experiments were performed with a minimum of 3 biological replicates. Statistical significance was determined by one-way or two-way analysis of variance (ANOVA) with comparisons to control as appropriate for each experiment, unless otherwise noted. Significance was assessed using Dunnett’s post-hoc multiple comparisons test. GraphPad PRISM v9 or higher and Statplus software package were used GraphPad Prism was used for all statistical analysis (SAS Institute: Cary, NC, USA) and data visualization. Error bars in all bar graphs represent standard error of the mean unless otherwise noted in the figure. No outliers were removed from the analyses. For all statistical analyses, a significant difference was accepted when P < 0.05.

## Acknowledgements

We would like to thank the patients and families for permission to publish this work. We thank Dr. Eric Payne and Dr. Suzanne Hoppins for providing the patient-derived fibroblast lines. We would also like to thank Anna Bright for technical assistance, Dr. Evan Krystofiak for providing EM expertise, Dr. Matthew Tyska, and Kari Seedle for providing advice and expertise with high-resolution microscopy. Dr. Kristopher Burkewitz, Dr. Jonathan Brunger, Dr. Kevin Ess, and Caleb Hayes for helpful scientific discussions about this study. Special thanks to Dr. J-P Cartailler and Dr. Shristi Shrestha, from Creative Data Solutions at Vanderbilt University, for their guidance on RNAseq data analysis. This work was supported by 1R35 GM128915-01 and 2R35GM128915-06 (VG), 1F31NS129296-01 (TB), and 1F31HD114431-01 (CB). All SIM and spinning disk confocal microscopy imaging and image analysis were performed in part using the Vanderbilt Cell Imaging Shared Resource, which is supported by NIH grants 1S10OD012324-01 and 1S10OD021630-01. The authors declare no competing financial interests.

## Author contributions

T. Baum and V. Gama conceived the study, designed experiments, interpreted data, and wrote the manuscript. V. Gama supervised and funded the research. T. Baum designed and carried out all the cell biology experiments, with technical assistance from C. Bodnya and J. Costanzo. All authors edited the document.

