## Supplemental Information for "Patient mutations in DRP1 perturb synaptic maturation of cortical neurons"

**Supplementary Figure Legends**

**Supplementary Figure 1 | Cortical differentiation timeline** iPSCs with DRP1 mutations differentiated in Knock Out Serum Replacement media from day 0 and gradually transitioned into N2 media over 11 days with addition of dual SMAD inhibitory small molecules SB-431521 and LDN-193189. After day 11, cells are transitioned into neural maintenance media for the duration of culture.

**Supplementary Figure 2 | Gene expression level analysis** (**A**) Distribution of gene expression levels of control, G32A, and R403C cultures. (**B**) Heat map of correlation analysis between mutant DRP1 cultures and control. (**C**) PCA plot clustering of 35 DIV and 65 DIV mutant DRP1 cultures and control. (**D**) Coexpression venn diagrams of 35 DIV (top panel) and 65 DIV (bottom panel) G32A and R403C cultures compared to control. (**E**) Histogram of number of differential genes for each comparison combination.

**Supplementary Figure 3 | Differential gene expression for 65 vs 35 DIV cultures** Volcano plots of DEX genes in 65 vs 35 DIV G32A and R403C cultures

**Supplementary Figure 4 | GO and KEGG plots** (**A**) top ten GO (left panel) and KEGG (right panel) enriched terms from control 65 vs 35 DIV cultures (**B**) top ten GO (left panel) and KEGG (right panel) enriched terms from G32A 65 vs 35 DIV cultures (**C**) top ten GO (left panel) and KEGG (right panel) enriched terms from G32A 65 vs 35 DIV cultures.

#### Supplementary Figure 1

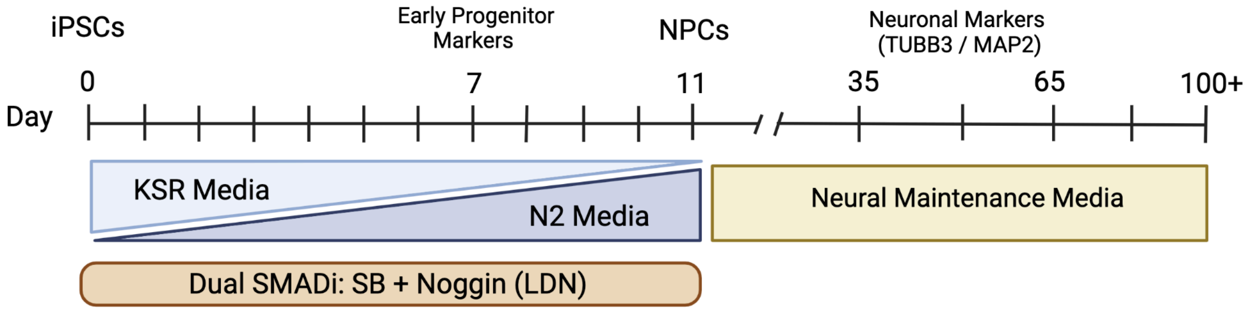

### Supplementary Figure 2

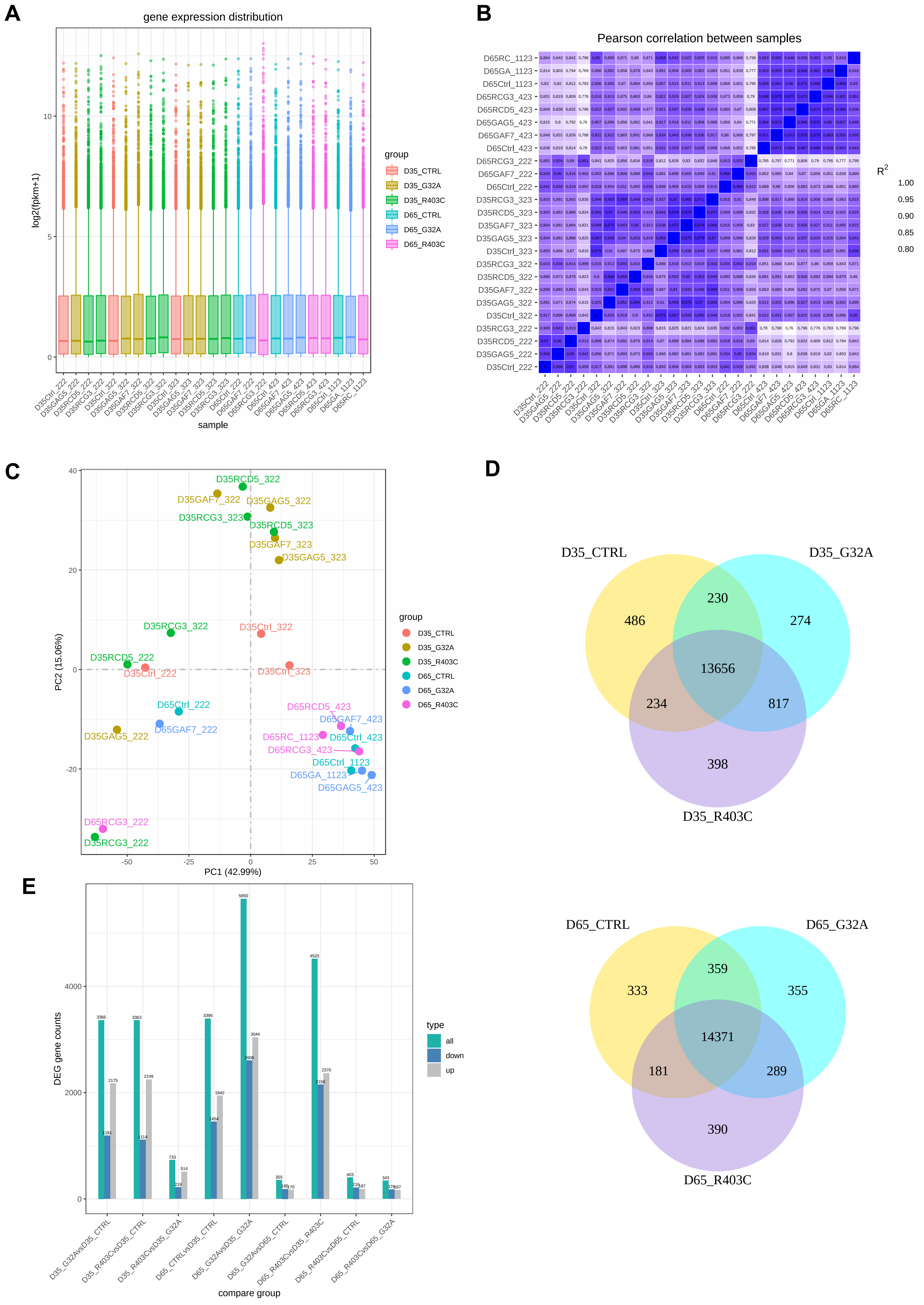

### Supplementary Figure 3

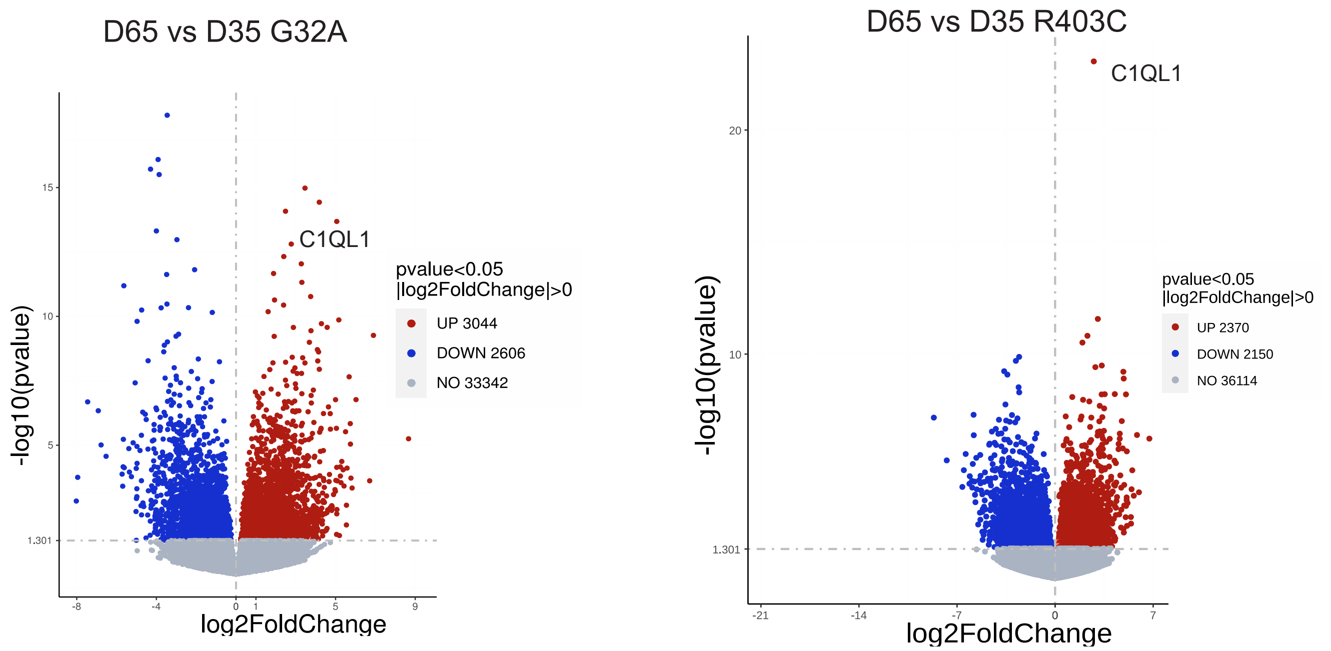

### Supplementary Figure 4

**A**

#### D65 vs D35 Control

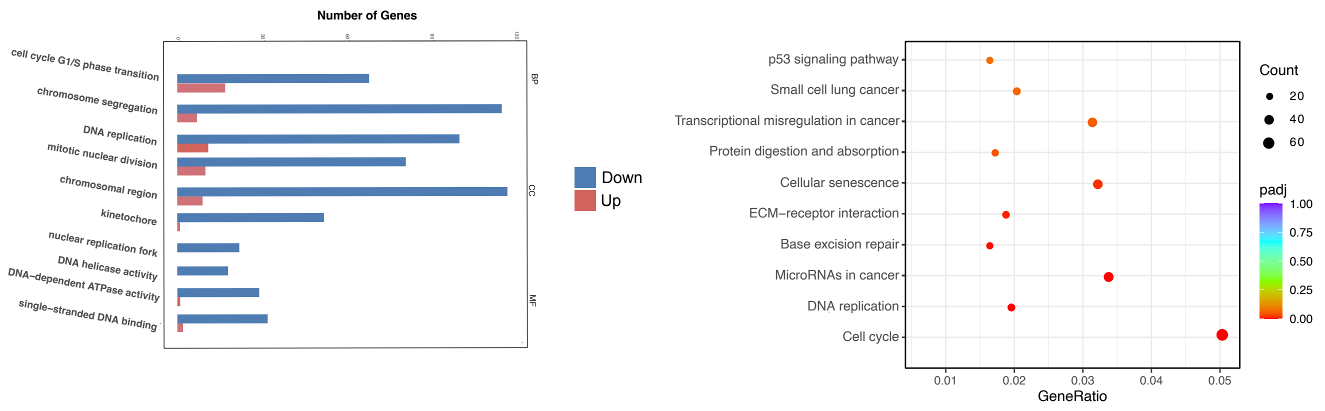


**B**

## D65 vs D35 G32A

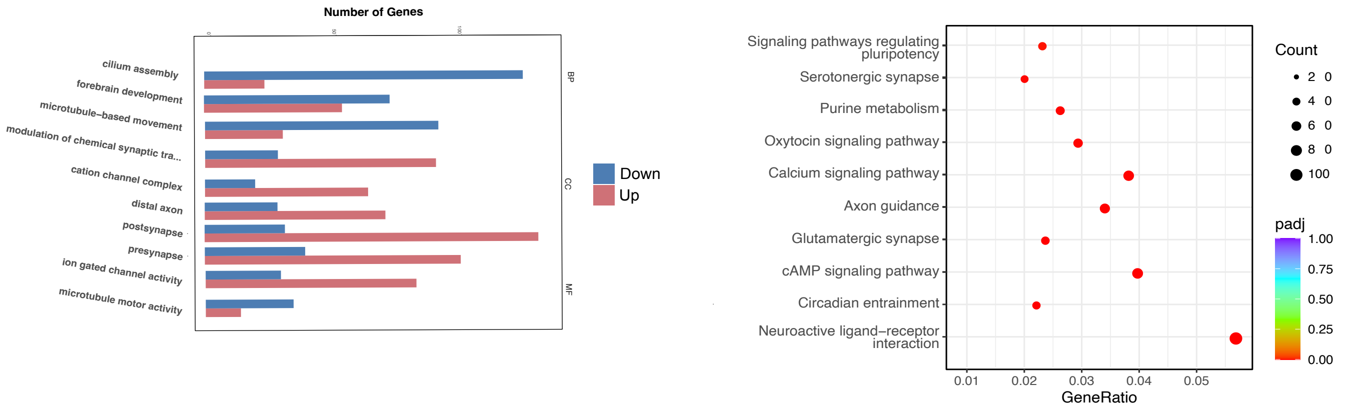


**C**

## D65 vs D35 R403C

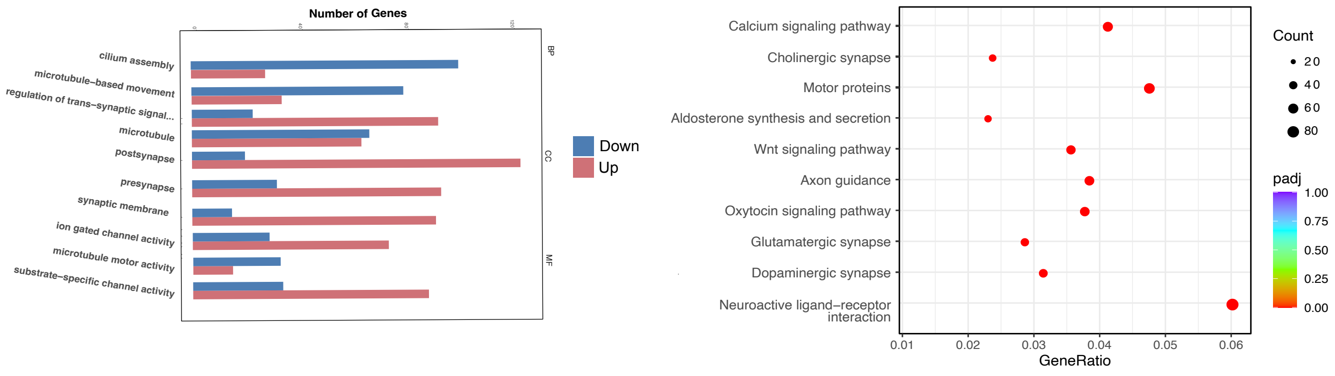
